# Bone marrow-on-a-chip: Emulating the human bone marrow

**DOI:** 10.1101/469528

**Authors:** Stefan Sieber, Annika Winter, Johanna Wachsmuth, Rhiannon David, Maria Stecklum, Lars Hellmeyer, Lorna Ewart, Uwe Marx, Roland Lauster, Mark Rosowski

## Abstract

Multipotent hematopoietic stem and progenitor cells HSPC reside in specialized stem cell niches within the bone marrow, that provide a suitable microenvironment for lifelong maintenance of the stem cells. Meaningful in vitro models recapitulating the in vivo stem cell niche biology can be employed for both basic research as well as for applied sciences and represent a powerful tool to reduce animal tests in preclinical studies. Recently we published the generation of an in vitro bone marrow niche model, capable of long-term cultivation of HSC based on an organ-on-a-chip platform. This study provides a detailed analysis of the 3D culture system including matrix environment analysis by SEM, transcriptome analysis and system intrinsic differentiation induction. Furthermore, the bone marrow on a chip model can serve to multiply and harvest HSPC, since repeated cell removal not compromised the functionality of the culture system. The prolongation of the culture time to 8 weeks demonstrate the capacity to apply the model in repeated drug testing experiments. The quality of the presented system is emphasized by the differentiation capacity of long-term cultivated HSPC in vitro and in vivo. Transplanted human HSPC migrated actively into the bone marrow of irradiated mice and contributed to the long-term reconstitution of the hematopoietic system after four and eight weeks of in vitro cultivation.

The introduced system offers a multitude of possible applications to address a broad spectrum of questions regarding HSPC, the corresponding bone marrow niche biology, and pathological aberrations.

## Introduction

In 1978, Schofield introduced the concept of a hematopoietic stem cell niche in the bone marrow capable of harboring native HSCs (Schofield, 1978). Since then, intensive research has revealed its crucial role in self-renewal, apoptosis, differentiation, quiescence, migration and immune privilege of HSCs (Morrison and Scadden, 2014). The HSC niche is a complex structure in the trabecular region of the bone. It is composed of multiple cell types, ECM and secreted factors which promote HSC maintenance, localization, and differentiation (Lilly et al., 2011). Various publications have pointed out the significance of direct cell-to-cell contact by ‘partner cells’ for the preservation of HSCs within their niche (Arai and Suda, 2007; Lilly et al., 2011). Distinct bone marrow stromal cells, besides endothelial cells, are assumed to be essential supporting cells for HSC preservation (Morrison and Scadden, 2014). Mesenchymal stromal cells (MSCs) expressing markers including CXCL12 and Nestin have been shown to provide factors that promote HSC maintenance (Yu and Scadden, 2016). Although the identification and characterization of the different MSC subtypes are still incomplete, various studies indicate that MSCs are an essential component for the maintenance of HSCs in the bone marrow niche (Yu and Scadden, 2016). The control of sustainment, mobilization and guided differentiation of HSPCs in the bone marrow niche has been a subject of considerable interest in recent times.

In our present day, animals are commonly used for the testing of emerging drugs. Apart from ethical considerations, this practice contains many downsides. Among these are the unreliable test results due to species-specific differences in rodent models. Concerning the bone marrow, these include among others the different set of surface markers being expressed on HSCs or the actual site of hematopoiesis which in human mostly takes place in the axial skeleton whereas all bones support hematopoiesis in mice (Yu and Scadden, 2016). It has been reported that less than 50% of animal tests predict human response accurately and 92% of new drugs, successful in animal test systems, fail in clinical trials (Hackam, D.G., and Redelmeier, 2006; Perel et al., 2007). Due to the high rate of failure in clinical trials and the associated high costs, the development of drugs with higher specificity and efficacy to treat various diseases is significantly delayed. To overcome this problem innovative *in vitro* 3D culture models are needed. In 2012, Sharma et al. presented a 3D hydrogel-based MSC HSPC co-culture system capable of keeping HSC in a quiescent state over seven days of culture. They attributed this to the formation of a hypoxia-gradient and interaction with the MSCs. Furthermore, Di Maggio et al. cultured freshly isolated nucleated cells of the human bone marrow on a hydroxyapatite scaffold in a perfused system for three weeks. Subsequently, HSPCs were successfully cultured in this scaffold for one week (Di Maggio et al., 2011; Sharma et al., 2012; Walasek et al., 2012). Concerning a bone marrow-on-a-chip, in 2014, Torisawa et al. introduced a bone marrow organoid which was engineered *in vivo* in a mouse employing murine hematopoietic stem cells. This construct was subsequently transferred to a microfluidic device and cultured for one week. (Kim et al., 2015; Torisawa et al., 2014). Unfortunately, no model was able to mimic the human hematopoietic stem cell niche meaning the continued sustainment of primitive HSCs while simultaneously allowing various hematopoietic populations to differentiate into their respective progeny.

In a preceding publication, we presented a proof-of-principle of a novel *in vitro* bone marrow model which utilizes a scaffold mimicking the structure and surface properties of the cancellous bone microstructure, thus, facilitating unobstructed interaction between the bone marrow-derived MSCs and umbilical cord-derived HSPCs (Sieber et al., 2018). The whole model is placed in a microfluidic device enabling the generation of different niches as well as the co-culture with other *in vitro* organ models. In this publication, by among others providing Illumina transcriptome analysis, long-term cultivation capacity and *in vivo* testing data, we are able to prove the functionality and robustness of this versatile pure *in vitro* bone marrow model, thereby exhibiting its vast potential. The here presented model demonstrates for the first time the successful long-term culture of functional multipotent HSCs in a dynamic environment for at least eight weeks. The model surpasses all previously presented 3D bone marrow models in culturing time of HSCs as well as in mimicking the *in vivo* environment (Kim et al., 2015). Predestining it as a model for sophisticated *in vitro* drug testing, thus, serving as an alternative to animal testing.

## Results

In this bone marrow model, we rebuild this environment in a hydroxyapatite-coated zirconium oxide scaffold with the intention of facilitating the long-term culture of primitive HSCs.

### Building a functional bone marrow model

The foundation for the successful culture of HSCs in the bone marrow model is laid by generating a suitable environment within the 3D hydroxyapatite coated zirconium oxide scaffold which mimics the situation observed *in vivo* (Fig.1A). MSCs isolated from bone marrow of the femoral head were cultured on a scaffold engineered with the intention of imitating the porous yet rigid properties of the human cancellous bone microstructure (Fig. 1B). After one week of culture the formation of an environment resembling the surroundings observed in the bone marrow *in vivo* was revealed (Sieber et al., 2018). A strong deposition of ECM was visible in the SEM overview pictures. At a higher magnification, the web-like network of ECM secreted by MSCs was observable. HSPCs could be seen to reside on the ECM covered surface of the ceramic. Furthermore, bridge structures that altered the architecture of the scaffold by spanning over cavities could be observed in various places throughout the ceramic. These bridge structures were also found in human bone marrow samples examined by our group (Fig. 1C and D). Subsequently, CD34^+^ HSPCs isolated from umbilical cord blood were cultured on the prepared ceramics within the MOC over the course of four weeks. HSPCs were extracted from the ceramic after 1, 2, 3 and 4 weeks of culture under dynamic conditions and stained for CD34 and CD38. Although the percentage of CD34^+^CD38^−^ HSPCs decreased over the time of culture, a substantial proportion of the regained cells, on average 32.96%, retained their primitive phenotype after four weeks of culture (Fig. 1E).

**Fig. 1:**
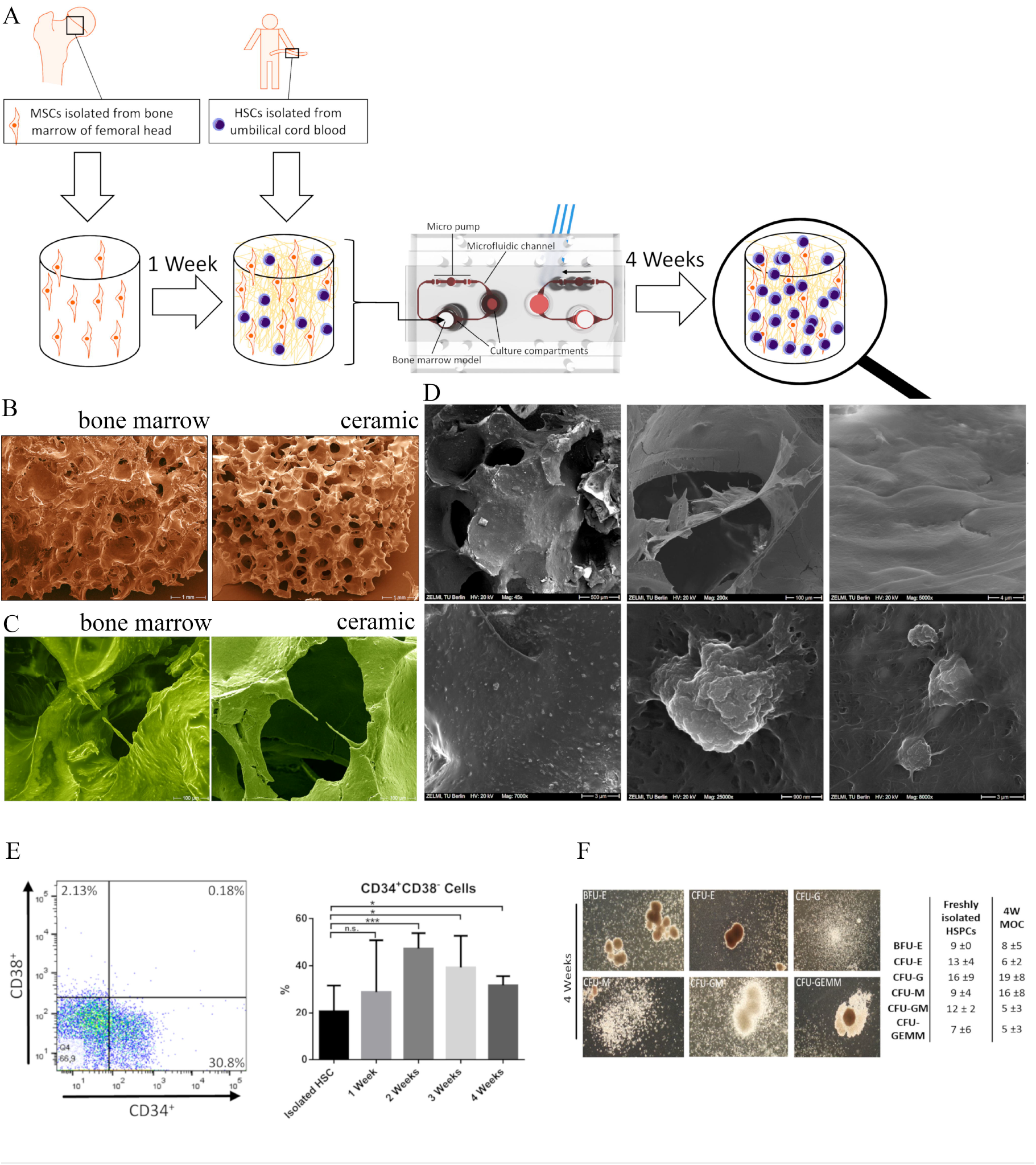
Hematopoietic stem cells are maintained for at least four weeks in the bone marrow model. **(A)** Bone marrow-derived MSC and umbilical cord blood-derived HSC were successively cultivated on the ceramic scaffold and subsequently integrated into the microfluidic organ on a chip platform. **(B)** The ceramic surrogate possesses high resemblance in structure and porosity to the in vivo counterpart **(C-D)** SEM analysis after one week of ceramic cultivation displayed the preparation of the HSC niche conditions. The MSC densely settled the ceramic, secreted extracellular matrix molecules and actively rearranged the ceramic cavities by bridge formation to provide a suitable niche microenvironment. **(E)** Representative FACS plot of HSPC extracted from the bone marrow model after 47 weeks of culture. Significant amounts of CD34^+^ single positive HSPC prove the capacity for long-term cultivation of hematopoietic stem cells (left) with even increase of the stem cells within the first three weeks of culture (right) (n=8). The elicited proliferative activity renders the system eligible for in vitro multiplication of the rare HSPC population. **(F)** The in vitro cultivated HSPC maintain their characteristic differentiation capacity in vitro with comparable efficacy to freshly isolated cells. The Table is showing the mean number of counted colonies of the CFU-GEMM assay performed with cells extracted after four weeks of culture (n = 7) or with freshly isolated HSPCs from umbilical cord blood.

To assess the conserved differentiation ability of the long-term cultured HSCs, a CFU-GEMM assay was conducted. Compared to freshly isolated HSPCs, the colony numbers were comparable to the freshly isolated HSPCs after four weeks of culture (Fig. 1F).

### Transcriptome analysis

To assess the impact of 3D cultivation and serum free-medium conditions on MSC transcriptional changes were determined by next-generation sequencing. Sets of 381 and 554 genes after one and four weeks respectively were defined to be differentially expressed with significant overlap. (Fig. 2B). Interestingly, clustered heatmap analysis of all expressed genes reveal the transcriptome properties of individual primary cells with donor intrinsic expression intensities characterized by donor-specific clustering (Fig. 2A). Comparative cluster analysis of regulated genes demonstrated a higher similarity of 3D cultivated cells compared to monolayer cells, but the primary cell character marked by individual transcription intensities was still evident within the ceramic grown cells (Fig. 2C). To assign affected biological processes and signal transduction pathways Gene Set Enrichment Analysis was performed. After the pre-cultivation phase to prepare HSC niche conditions mainly GO-terms connected to proliferative activity, cell cycle regulation and cytoskeleton rearrangement were overrepresented (Fig. 3A). According to the intensified cell-cell contact in 3D culture genes mediating focal adhesion, gap junction formation, and ECM-receptor interaction were determined to be overrepresented within the set of differentially expressed genes (Fig. 3A). To assess the quality of cell division regulation upon ceramic culture individual cluster analysis for the term `GO:0007049` “cell cycle regulation” was performed. As indicated for the entire set of regulated genes, the majority of the cell cycle genes were downregulated with a similar pattern of sample clustering (Fig. 3B). Cell cycle progression is regulated by cyclin-dependent kinases (CDKs) and their activation by members of the cyclin (CCNs) family or the inhibition by CDK inhibitory molecules (CDKNs). Cluster analysis with the focus on this three protein families revealed downregulation of supportive molecules and upregulation of the inhibitors of cell cycle progression within the set of differentially expressed genes (Fig. 3C).

**Fig. 2:**
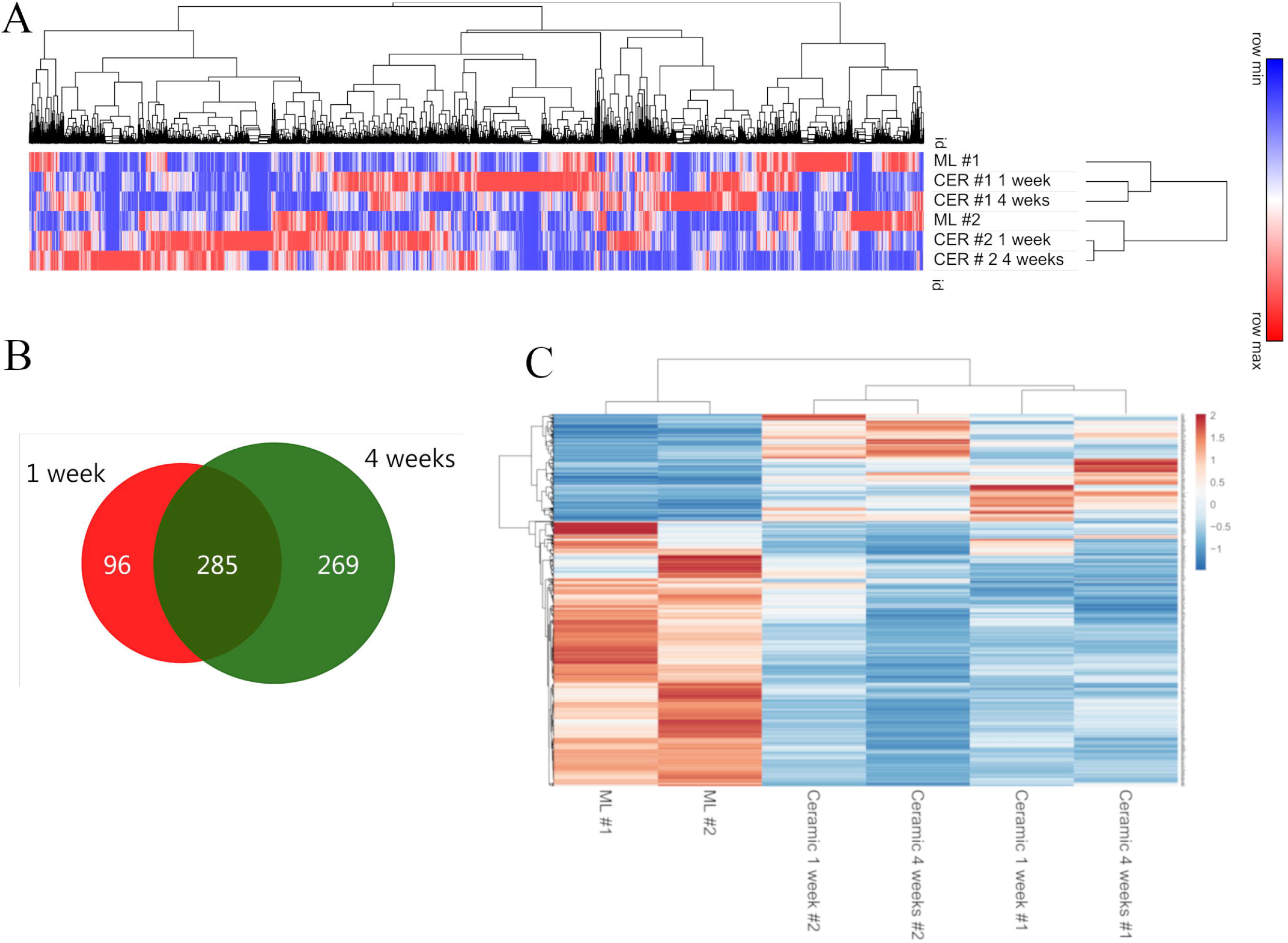
Fig. 2: Comprehensive transcriptome analysis by next-generation sequencing **(A)** After one and four weeks of scaffold cultivation of MSCs RNA was isolated and differentially expressed genes were determined by sequencing technology. Cluster analysis of all expressed genes revealed a higher resemblance of 3D cultivated cells compared to monolayer MSCs. However, the expression pattern and the cluster formation is dominated by the primary cell character marked by individual intrinsic expression strengths (n=2). **(B)** An increasing set of 381 and 554 differentially expressed genes with significant overlap after one and four weeks respectively account for a progressive process of MSC alteration upon ceramic cultivation [the DESEQ2 algorithm; nominal p-value < 0,05; FD>2) provided by DE analysis (https://yanli.shinyapps.io/DEApp/). **(C)** Clustered heat map analysis with a focus on differentially expressed genes substantiates the resemblance of 3D cultivated cells, however, within the clustered group of MSCs on ceramic the primary cell character of individual donors marked by respective transcription intensities remains evident.

**Fig. 3.**
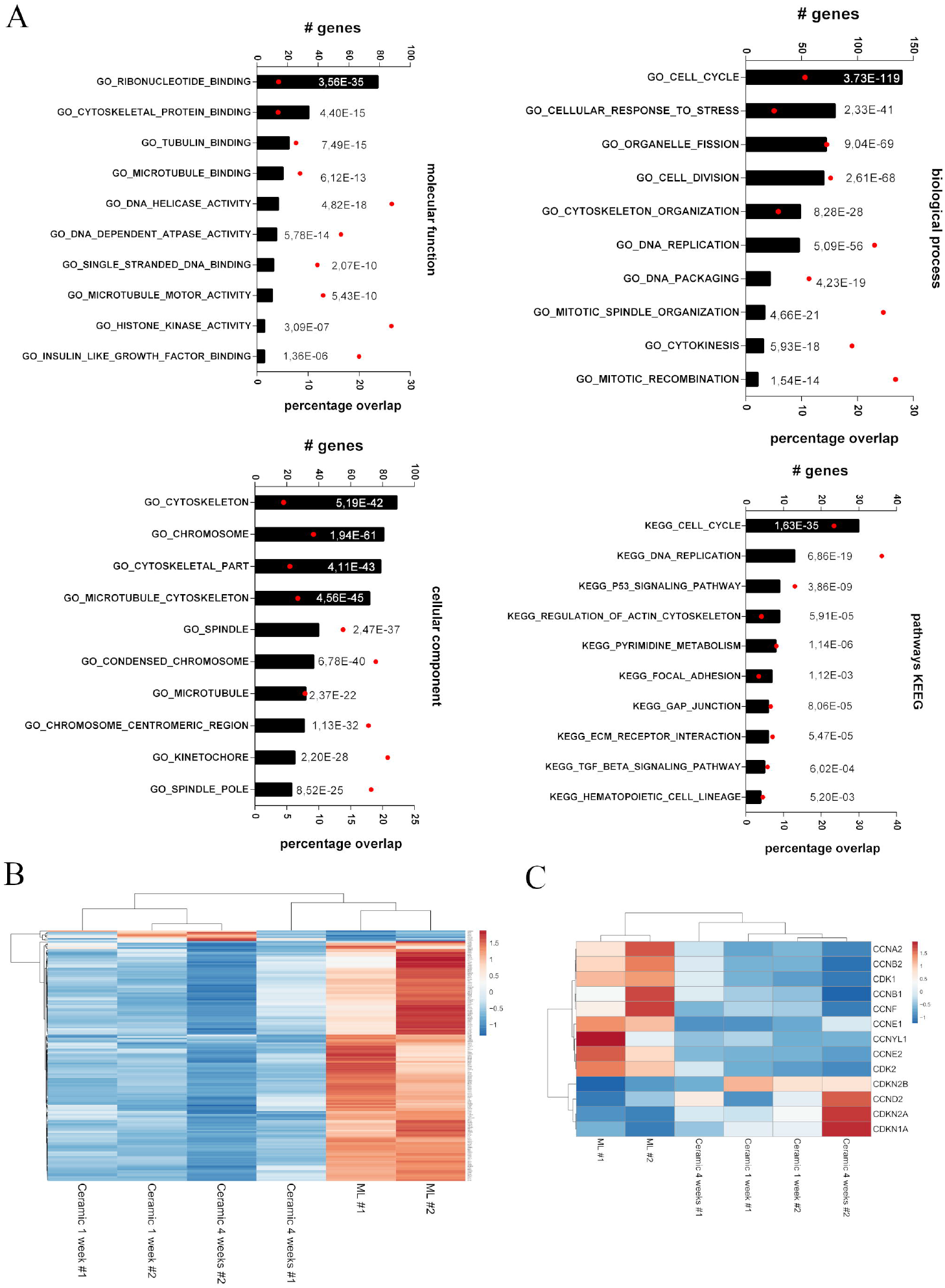
Gene set enrichment analysis of differentially expressed genes designated the cell cycle and related pathways to be the most prominent affected biological function [http://software.broadinstitute.org/gsea/index.jsp; broad institute, Cambridge] **(A)** Differentially expressed genes were analysed for overrepresented GO-terms regarding molecular function, biological process, cellular component, and KEGG pathways. Cell cycle regulation and related terms, cytoskeleton organization and cell to cell or cell to matrix interaction are highlighted to be the most influenced functions [bares - the number of differentially expressed genes in the corresponding GO_term; red dot percentage overlap of differentially expressed genes within the group; p-values for each GO-term are given in numbers). **(B)** Heatmap cluster analysis of differentially expressed gene of GO-0007049 (#cell cycle regulation) displayed downmodulation of the majority of genes with similar cluster properties observed for the entire set of regulated genes. **(C)** Focused cluster analysis of the central regulators of the cell cycle progression revealed upregulation of cyclin-dependent kinase inhibitors (CDKN) and downregulation of CDKs and the activating cyclin molecules. 3D cultivation of MSC downregulates the cell cycle and cell proliferation.

Interestingly, GSEA derived no GO terms related to stem cell maintenance, stem cell niche or bone marrow microenvironment. To define the functionality of the generated artificial HSC niche, the expression of molecules known to be essential for HSC maintenance were determined. The majority of the analyzed molecules are expressed by MSC independent of the cultivation method with no significant regulation upon 3D culture (Fig. 4 A-C). Only a very few genes like thrombopoietin CASR or delta-like molecules (DLL) are very little or not expressed at all. In reference to DESEQ2 analysis, only a small number of genes were differentially expressed. BMP-2 and Angiopoietin-1 displayed elevated expression whereby molecules like JAG1 and KITLG showed even decreased RNA levels upon ceramic cultivation Fig. 4 A and B).

**Fig 4.**
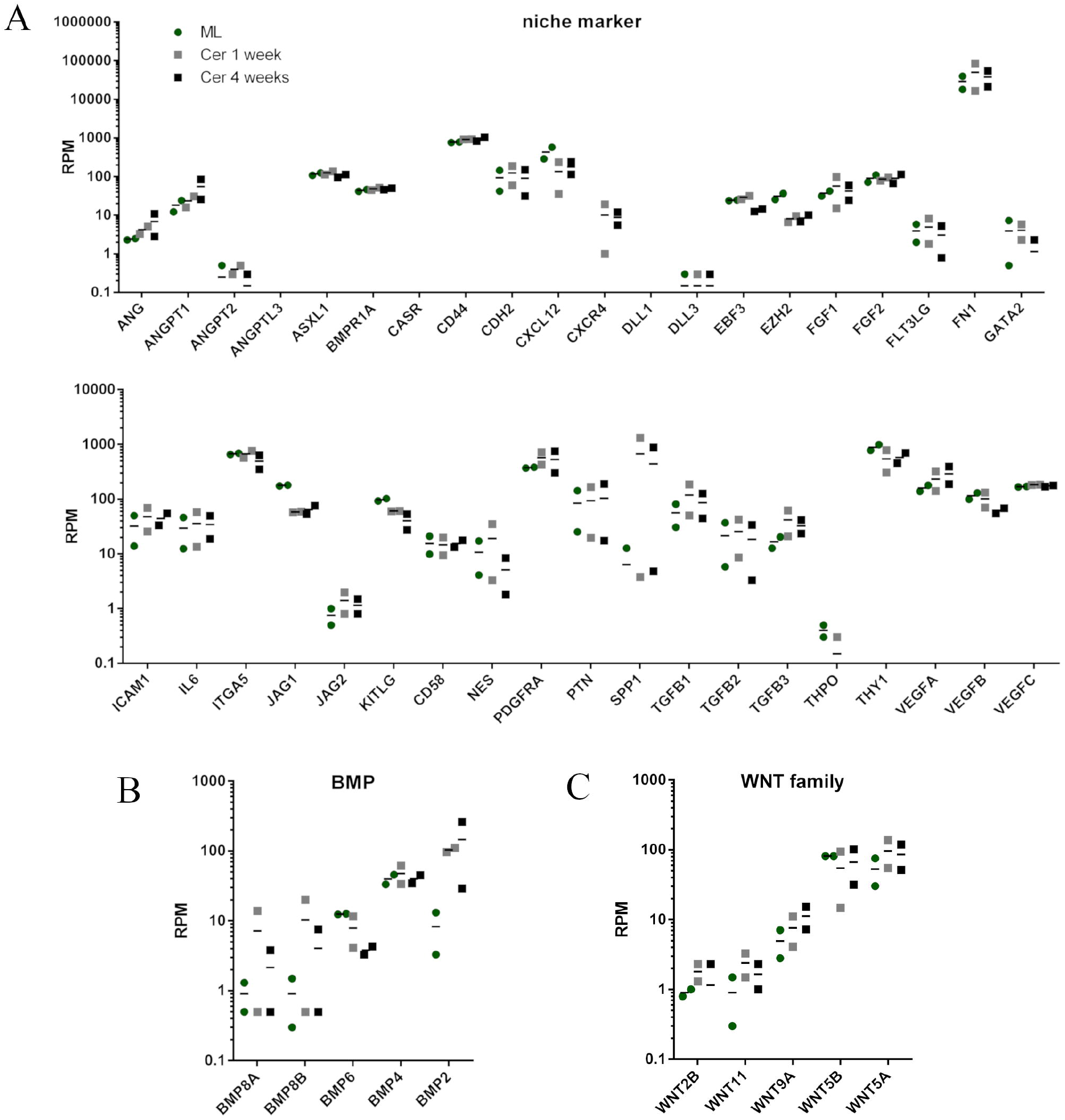
The majority of genes known to affect the HSC niche biology are expressed independently from the cultivation condition. **(A)** RPM gene expression values from next-generation sequencing for genes involved in HSC maintenance in the bone marrow and with focus on BMPs **(B)** and the WNT family **(C)** are shown. The genes are not differentially expressed but present in significant strength to fulfill their functions in the in vitro cultivation system independent of induction upon the ceramic cultivation.

### HSPCs differentiate into various lineages

During the culture of the HSPCs on the MOC, cells that appeared to be HSPCs were observed in the medium reservoir, i.e., the culture compartment not occupied by the ceramic. To investigate if the cells had started to differentiate after they had left the environment of the ceramic, the ceramic and circulating HSPCs were analyzed separately.

Native HSCs (CD45RA^−^CD34^+^CD38^−^CD90^+^) resided more abundantly in the artificial bone marrow niche in the ceramic compared to the circulation. After 2, 3 and 4 weeks of culture, a significant difference between the two compartments could be detected (Fig. 5A). The difference was also apparent when looking at the absolute cell numbers, although only the difference after three weeks of culture was significant (Fig. 5D). In line with these observations, more cells expressing the myeloid marker CD45RA were found in the circulation. In percentage as well as absolute cell numbers, a significant difference could be observed after three weeks of culture (Fig. 5B and E). Although there was no significant difference in the other weeks, a clear trend was evident. The same applied for the expression of the erythroid marker CD36.

**Fig. 5:**
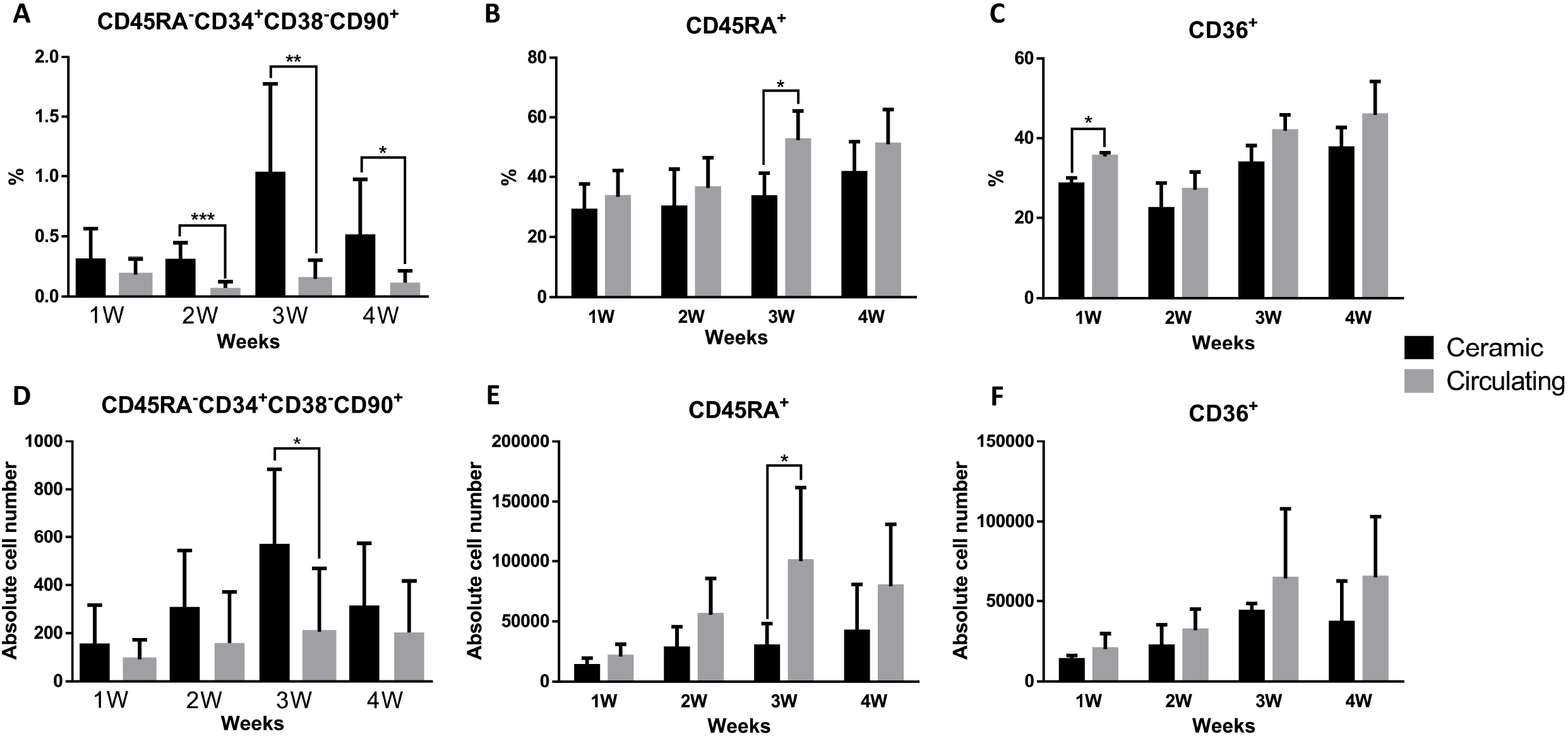
Cells extracted from the ceramic and the circulation display different surface marker expression patterns. **(A-C)** Percentage and **(D-F)** absolute cell numbers of CD45RA−CD34+CD38−CD90+, CD45RA+ or CD36+ cells cultured on the MOC for 1, 2, 3 and 4 weeks. Cells extracted from the ceramic and the circulation were stained and analyzed separately by flow cytometry. **(A and D)** HSCs with the phenotype CD45RA^−^CD34^+^CD38^−^CD90^+^ are predominantly located in the ceramic scaffold with a significant difference after three weeks, while the more differentiated progeny cells expressing the surface markers **(B and E)** CD45 and **(C and F)** CD36 appear enriched in the circulating system. Asterisks indicate a significant difference between the two values. Error bars represent the standard deviation (n=5).

Having established that a significant difference between the cells extracted from the ceramic and the circulation exists, a more extensive analysis of the subpopulation of differentiated HSPCs was carried out. Due to a two-week stabilization phase, the cultures lasted for 3, 4 and 5 weeks. The stabilization phase was introduced to generate a higher starting population. For the characterization of the cells potentially differentiating in the neutrophil lineage, the occurrence of GMPs, myeloblasts, myelocytes, and neutrophils was investigated. FACS analysis applying the individual marker molecules CD34, CD38, CD45RA, CD36, CD15 and CD16 and their combinations for the distinct differentiation stages were conducted. GMPs and myelocytes were the cell types essentially found whereas myeloblasts and neutrophils were barely present in both compartments, the ceramic and the circulation system (Fig. 6).

**Fig. 6.**
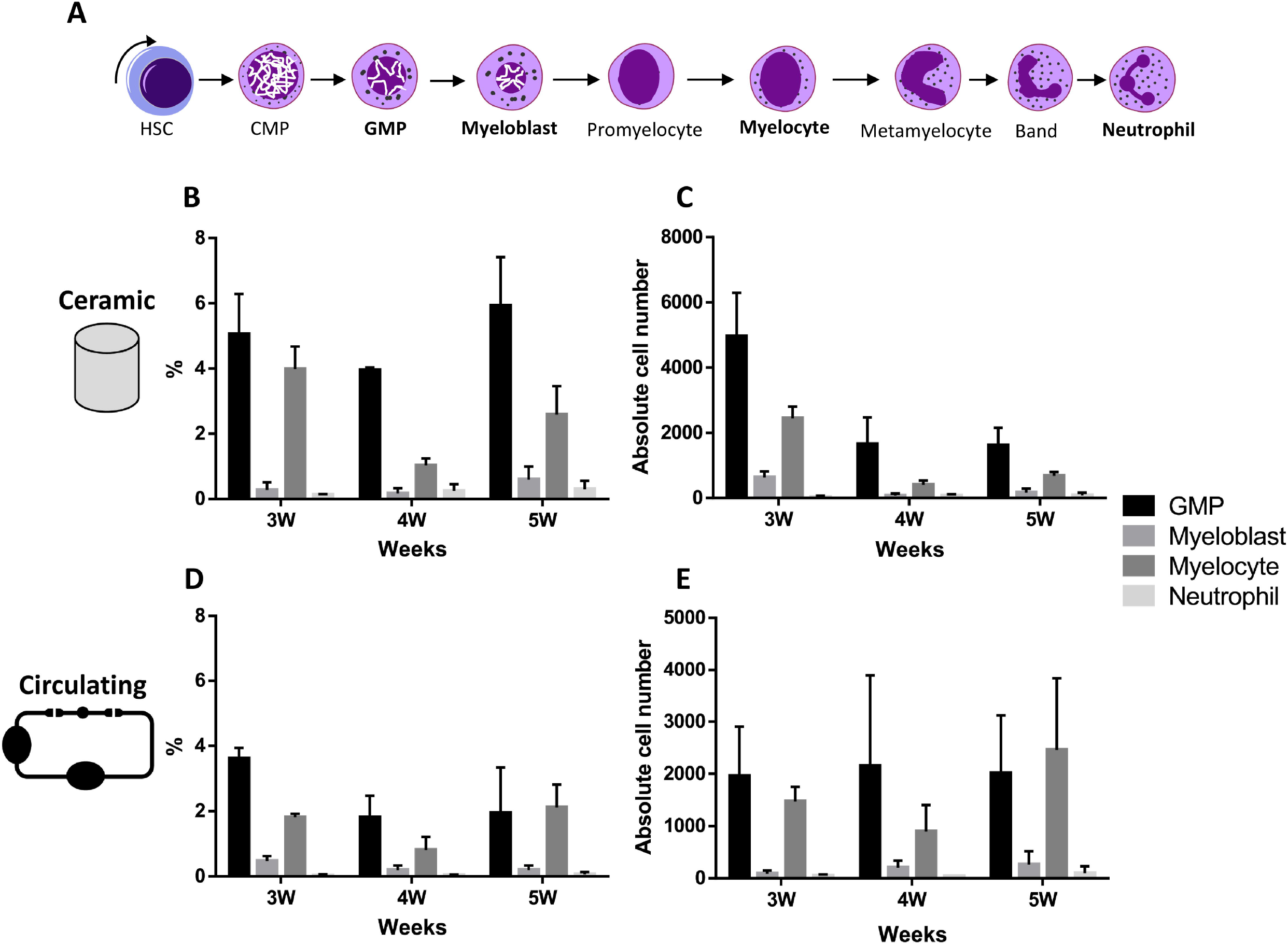
Differentiation of HSPCs towards the neutrophil lineage is observable. **(A)** Scheme of Neutrophil differentiation stages of HSPCs. Cells were extracted to the designated time points from the ceramic compartment or the circulating periphery. For the characterization of the cells potentially differentiating in the neutrophil lineage, the occurrence of GMPs (CD36^−^CD45RA^+^CD34^+^CD38^+^CD15^−^CD16^−^), myeloblasts (CD36^−^CD45RA^+^CD34^+^CD38^+^CD15^+^CD16^−^), myelocytes (CD36^−^CD45RA^−^CD34^−^CD38^+^CD15^+^CD16^−^) and neutrophils (CD36^−^CD45RA^−^CD34^−^CD38^+^CD15^+^CD16^+^) were investigated (Attar, 2014). The cell types investigated in the experiment are written in bold. **(B-E)**. Percentage and absolute cell number of GMPs, myeloblasts, myelocytes, and neutrophils extracted from the ceramic or the circulation after 3, 4 and 5 weeks of culture, respectively. Error bars represent the standard deviation (n=3). The differentiation pattern depicted in (A) was modeled after Stiene-Martin et al. Stiene-Martin, A. Leukocyte Development, Kinetics, and Functions. clinicalgate (2015).

### The bone marrow model is robust

The cultivation system is characterized by augmentation of HSC within the first weeks of culture. Therefore, the option to harvest cells from the system during a running experiment for multiple applications was assessed. Furthermore, a positive result opens up the possibility to continually monitor various parameters in bone marrow safety studies without ending the experiment. After a two-week stabilization phase, all, half or none of the HSPCs extracted from the circulation during medium exchange were returned to the MOC. After this stabilization period, the culture ran for an additional 1, 2 or 3 weeks.

Regarding the CD34^+^CD38^−^ cells, more cells were found in the circulation compared to the ceramic after three weeks of culture after complete cell reintroduction confirming the accumulation of HSC in early culture phases. After week five, the percentage of CD34^+^CD38^−^ cells appeared to be evenly distributed except for the 0% reintroduction group where a higher rate was seen in the circulation. The impact of cell removal was undoubtedly visible but not significant. Except for the cells extracted from the ceramic after five weeks of culture, a clear gradation was observable between the three different groups (Fig. 7).

**Fig 7:**
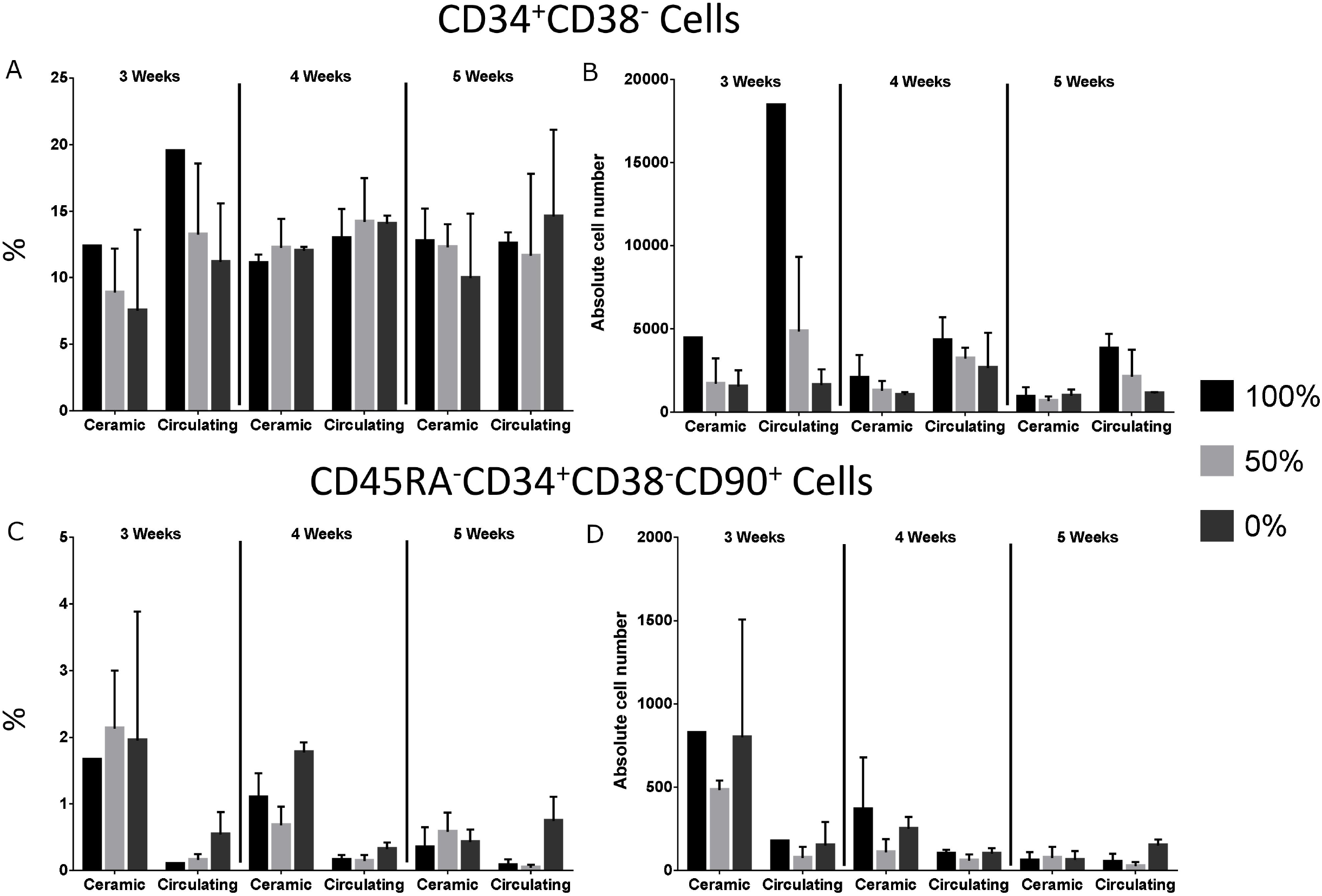
Impact on the hematopoietic population by reintroducing 100%, 50% or 0% of cells during medium exchange demonstrates the robustness of the system. **(A-B)** Percentage and absolute cell numbers of CD34+CD38− cells and **(C-D)** of CD45RA−CD34+CD38−CD90+ cells collected from the ceramic or the circulation after 3, 4 or 5 weeks of culture. During the medium exchange, 100%, 50% or 0% of the cells were reintroduced into the MOC. The removal of the cells showed only small effects suggesting an intrinsic self-stabilization capacity of the cultivation system. A two-week long stabilization phase with 100% reintroduction of cells preceded the experiment. Error bars represent the standard deviation.

Native HSCs (CD45RA^−^CD34^+^CD38^−^CD90^+^) were more abundantly identified in the ceramic as shown in Figure 5 and 7. An exception was the absolute cell number detected after five weeks of culture which were evenly distributed. A gradation as seen for the CD34^+^CD38^−^ cells was not observable. Interestingly, the number of native HSCs detected for the 0% reintroduction group was higher than the 50% reintroduction group and comparable to the 100% reintroduction group suggesting a renewal of the HSC subpopulation upon cell removal (Fig. 7).

### Culture of the bone marrow model on the MOC for eight weeks

After successfully culturing the bone marrow model on the MOC for four weeks (Sieber et al., 2018), the culture time was extended to eight weeks. A longer culture time presents the possibility to perform long-term repeated drug tests.

For the various populations, a variance between the measured percentage and the regarding absolute cell number was observable. While the percentage of CD34^+^CD38^−^ cells decreased, the absolute cell numbers remained stable between 4 and 8 weeks of culture (Fig. 8 C and H). The CD45RA^−^CD34^+^CD38^−^**CD49f**^+^ and CD45RA^−^CD34^+^CD38^−^**CD90**^+^ populations nearly vanished after eight weeks of culture while the CD45RA^−^CD34^+^CD38^−^**CD133**^+^ population only declined regarding the percentage (Fig. 8F and K).

**Fig. 8.**
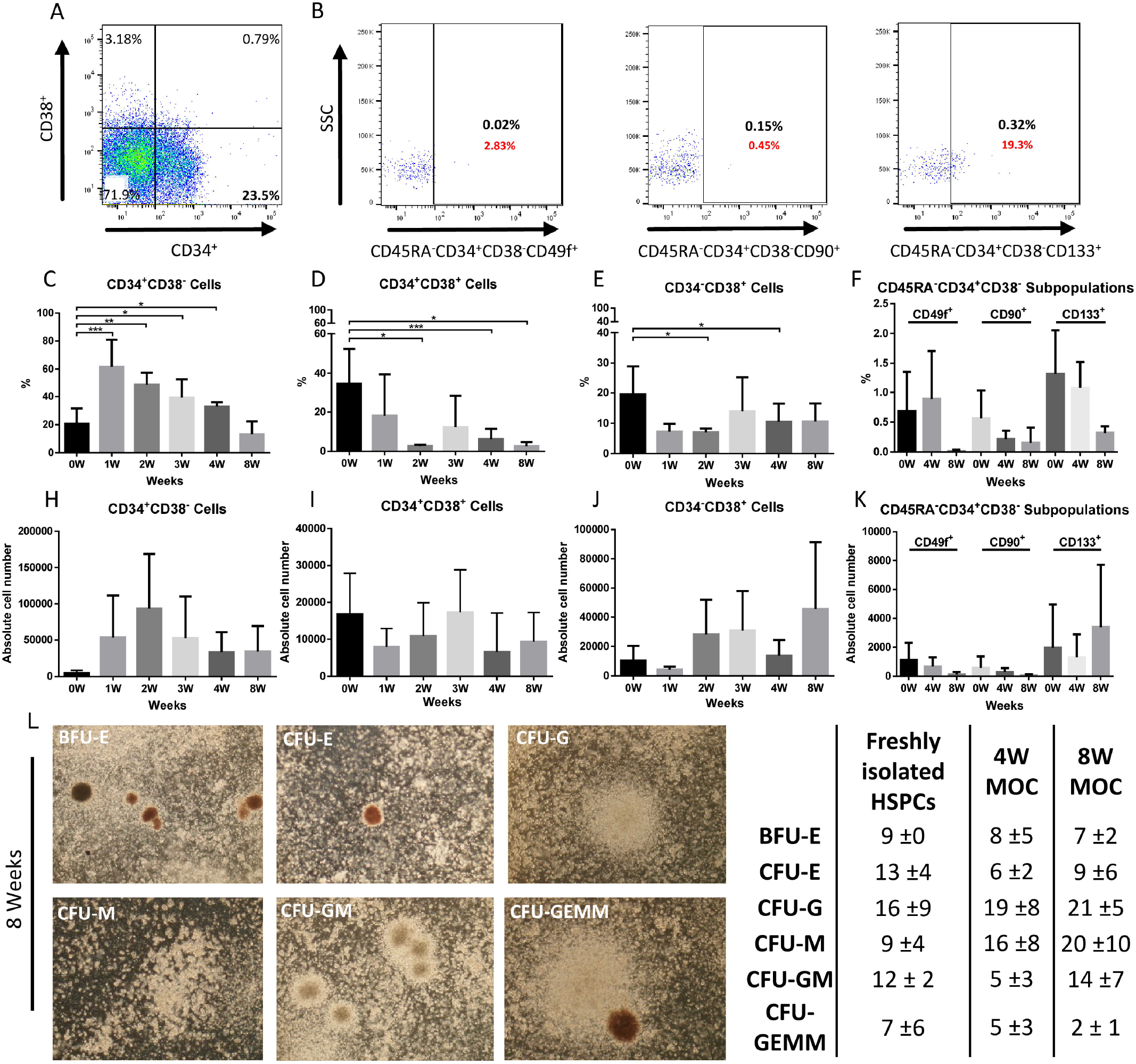
HSCs maintain their primitive phenotype in dynamic conditions over the course of eight weeks **(A)** Representative FACS plots of HSPCs extracted from the ceramic after eight weeks of culture on the MOC. Cells were stained for CD34 and CD38. **(B)** Representative FACS plots of CD45RA−CD34+CD38−CD49f+, CD45RA−CD34+CD38−CD90+ and CD45RA−CD34+CD38−CD133+ cells. The black numbers refer to the percentage of positive cells gated in the FSC/SSC, whereas the smaller, red numbers refer to the percentage of positive cells of the parent gate. **(C-E)** Percentage of CD34+CD38−, CD34+CD38+ and CD34−CD38+ cells over the course of eight weeks of culture in comparison to freshly isolated HSPCs. **(F)** Percentage of CD45RA−CD34+CD38−CD49f+, CD45RA−CD34+CD38−CD90+ and CD45RA−CD34+CD38−CD133+ cells directly after isolation, four weeks or eight weeks of culture. **(H-K)** The corresponding absolute cell numbers to **C-F**. The bridge connecting two values with the asterisk represents a significant difference. The error bars represent the standard deviation (n=3) **(L)** HSPCs are still able to differentiate after eight weeks of culture. Representative pictures of burst-forming-unit erythrocyte (BFU-E), colony-forming-unit erythrocyte (CFU-E), granulocyte (CFU-G), macrophage (CFU-M), granulocyte, macrophage (CFU-GM) and granulocyte, erythrocyte, macrophage, megakaryocyte (CFU-GEMM) colonies. The colonies originated from cells that were cultured under dynamic conditions for eight weeks (n=3). Table shows mean values and standard deviation of counted colonies of the CFU-GEMM assay performed with cells extracted after 4 and 8 weeks of culture in dynamic conditions or with freshly isolated HSPCs from umbilical cord blood.

By performing a CFU-GEMM assay, we could test whether the HSCs had kept their ability to differentiate *in vitro*). No substantial differences could be observed in comparison to the other culture times. (Fig. 8L)

### HSCs cultured in the bone marrow model engraft in irradiated immunocompromised mice

Transplantation of cells into irradiated immunocompromised mice is the gold standard assay to determine the stem cell state of hematopoietic stem cells. It demonstrates that HSCs are still able to long-term repopulate a vacant bone marrow niche and are therefore native stem cells.

The quantification of human CD45 expressing cells in the blood samples taken four, eight and twelve weeks after the injection of 4 weeks in vitro cultured HSPCs into recipient mice revealed an approximation of the values to the control of freshly isolated cells. No significant difference was detectable between the two values 12 weeks post-injection (Fig. 9 A-C). The same applied to the B cell (CD45^+^CD19^+^), monocyte (CD45^+^CD14^+^) and NK cell (CD45^+^CD56^+^CD16^+^) populations within human CD45^+^ cell compartment. The only exception was the occurrence of human T cells (CD45^+^CD3^+^) after 12 weeks which were significantly lower than the control within the human cell subset (Fig. 9D). All percentages, except for leukocytes, refer to the CD45^+^ gate.

**Fig. 9.**
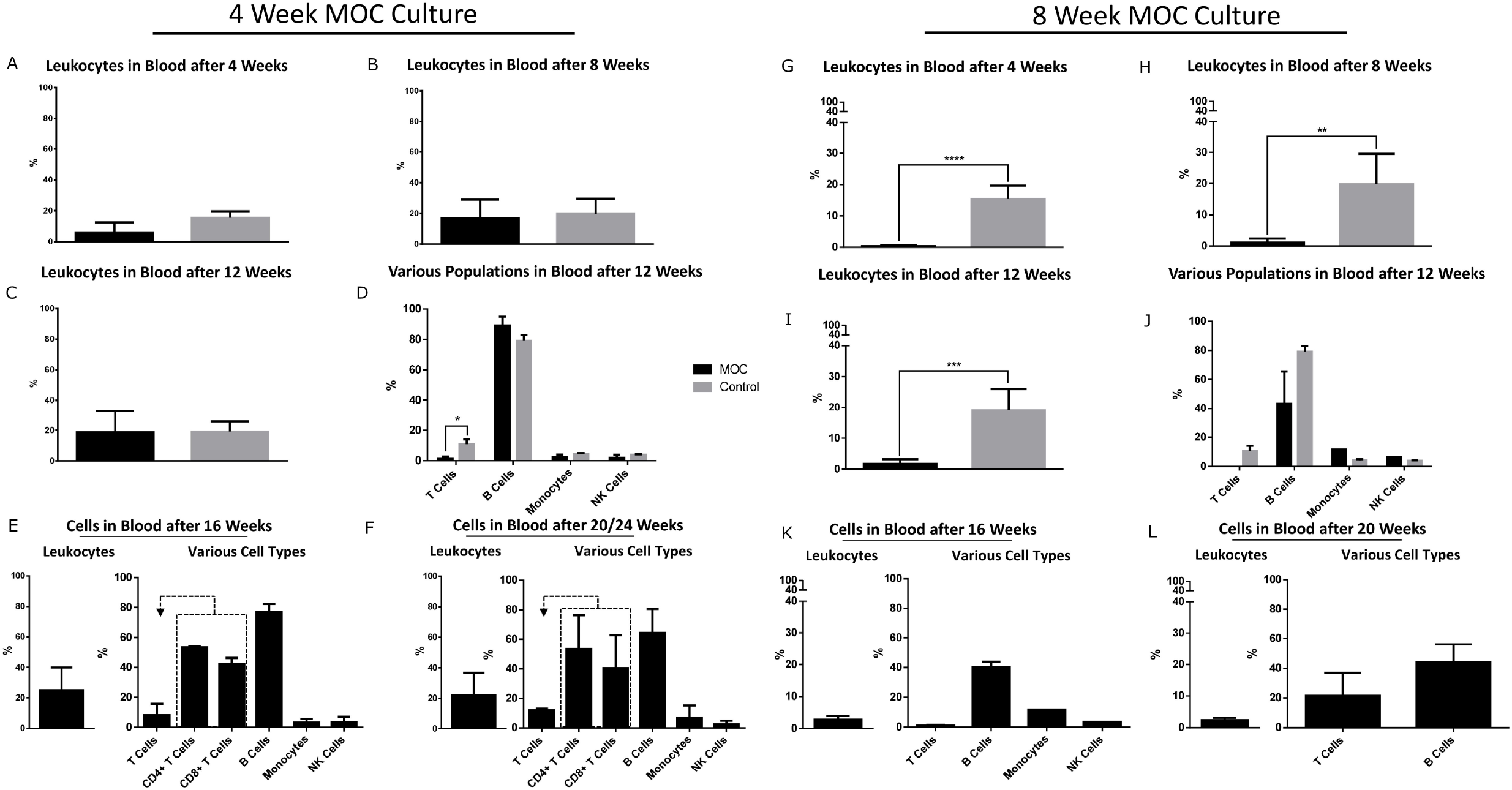
HSCs cultured for four weeks and eight weeks on the MOC can engraft in irradiated immunocompromised mice after transplantation. HSPCs freshly isolated from umbilical cord blood or cultured for four or eight weeks on the MOC were injected into irradiated immunocompromised mice. Blood samples were taken to the indicated time points. **(A-C)** Percentage of human CD45^+^ cells detected in the blood samples taken 4, 8 and 12 weeks after the injection of the HSPCs. **(D-F)** Analysis of particular immune cell populations (T cells, B cells, Monocytes, and NK) of human origin after 12 weeks **(D)**, 16 weeks **(E)** and 20 weeks **(F)** of culture. The percentages of CD4^+^ and CD8^+^ T cells refer to the T cell gate as indicated by the dotted rectangle. **(G-L)** Transplantation experiments were repeated with eight weeks in vitro cultivated HSPCs with the equivalent evaluation approach. **(G-I)** Percentage of human CD45^+^ cells detected in blood samples taken 4, 8 and 12 weeks after the transplantation. **(J-L)** Evaluation of individual human immune cell populations **(J)** 12 weeks, **(K)** 16 weeks and **(L)** 20 weeks after cell administration. As in figure **E and F**, the percentages of CD4^+^ and CD8^+^ T cells refer to the T cell gate as indicated by the dotted rectangle. The error bars represent the standard deviation.

After 16 weeks, the number of human leukocytes, T cells, monocytes, and NK cells had increased further. The B cell population did decrease insignificantly. No control was available for this time point (Fig 9 E). After 20 weeks the various populations stayed on the same level except for T cells which had further increased (Fig. 9F).

In a second *in vivo* experiment, HSPCs were cultured for eight weeks in the bone marrow model on the MOC before they were injected into an irradiated immunocompromised mouse. The same controls used in the experiment above were employed.

Compared to the first experiment, the number of human CD45 expressing cells was significantly lower compared to the controls after 4, 8 and 12 weeks (Fig. 9G - I). Additionally, no T cells could be observed after 12 weeks, whereas B cells, monocytes, and NK cells were present in a similar quantity as found in control experiments (Fig. 9J). After 16 and 20 weeks, the number of human leukocytes in the blood had further increased, which especially applied to the number of T cells measured (Fig. 9K-L).

## Discussion

The general aim was the establishment of a model capable of mimicking the human *in vivo* bone marrow environment. The model should be capable of sustaining native HSCs for at least eight weeks. The HSC bone marrow niche is a complex environment involving different cell types, their various ways of interaction, and specific physical microenvironments (Crane et al., 2017). This certain surrounding could only be reconstructed in a three-dimensional platform.

### Generating a bone marrow model

The foundation for the successful culture of HSPCs in the bone marrow model was laid by creating a suitable environment which mimics the situation observed *in vivo*. Thus, primary bone marrow-derived MSCs were employed and cultured on the hydroxyapatite-coated Sponceram 3D ceramic scaffold. This scaffold was chosen for its human bone marrow mimicking properties. The pore size and structure of the scaffold is comparable to the human bone marrow of the femoral head while hydroxyapatite is a close analog of bone apatite, found in hard tissue in all vertebrates (Junqueira, 2003; Murphy et al., 2010).

After four weeks of culture in the serum-free HSPC medium, SEM images demonstrated that the niche was still densely populated and secretion of ECM could be observed. Interestingly, bridge structures that altered the architecture of the scaffold by spanning over cavities could be seen in various places throughout the ceramic. These bridge structures were also observed in human bone marrow samples examined by our group. Thus, the MSC actively change the environment to their needs and expand the surface by bridging the cavities. The deposition of ECM molecules may contribute to the niche biology since essential growth factors and other essential signaling molecules secreted by the stromal cell compartment can be stored in the rearranged structures.

After the addition of the CD34^+^ HSPCs isolated from umbilical cord blood, the bone marrow model was cultured for up to eight weeks. During culture, the HSCs retained their native phenotype. The reduction of CD34^+^CD38^−^ and CD45RA^−^CD34^+^CD38^−^**CD90**^+^ HSPCs over the course of the culture can very likely be ascribed to part of the population starting to differentiate. The decline of the latter population indicates the reduced capability to repopulate a vacant bone marrow niche. Nevertheless, the population was still present and approximated the *in vivo* situation regarding the abundance, where HSC present a very rare population with an estimated frequency of 0.01% of total nucleated cells (Brendel et al., 2014; Gullo et al., 2015; Notta, F., Doulatov, S., Laurenti, E., Poeppl, A., Jurisica, I., and Dick, 2011; Rossi et al., 2011). The assumption of reduction due to cell differentiation is strengthened by the increase of the total cell number of cells positive for the differentiation markers CD45RA or CD36 as well as the elevated cell number of the CD34^+^CD38^+^ and CD34^−^CD38^+^ populations. An explanation might be given by considering the different environments depicted in the SEM pictures, cells appearing to be HSPCs were found embedded in ECM or residing directly on the hydroxyapatite-coated surface of the scaffold. Additionally, direct interactions between HSPCs and MSCs, as well as HSPCs and other HSPCs, were detected (data not shown). These different culture environments within the scaffold could explain the diverse HSPC populations identified. A direct interaction either with nestin-expressing MSCs or the embedding in ECM could be beneficial in preserving the native HSC phenotype. In contrast, sole HSPCs residing on the hydroxyapatite surface might be prone to differentiation (Morrison and Scadden, 2014). The CFU-GEMM assay showed that the cells exhibited a preserved myeloid differentiation capability of the long-term cultured HSPCs.

### 3D cultivation mainly affects proliferate activity

Interestingly, 3D cultivation of MSC rather influenced the proliferative activity of the cells than the induction of HSC maintaining molecules. Transcriptome analysis demonstrated the down-regulation of the cell cycle and related cellular processes like DNA replication. Molecules known to be essential for HSC maintenance in the bone marrow niche are expressed independently from 2D or 3D culture condition (Crane et al., 2017; Pinho et al., 2013). This finding is in accordance that MSC can be used as a feeder layer to cultivate and to expand CD34^+^ HSC in 2D cultivation systems (Magin et al., 2009; Michalicka et al., 2017). However, in these studies, the cultivation time does not exceed a time frame of 14 days. In our system long-term cultivation of native HSC at least up to 8 weeks is feasible. The maintenance of native HSC in vitro requires serum-free culture condition. One beneficial effect might be attributed to MSC survival by proliferation inhibition accompanied by concurrent maintenance of HSC niche factor expression upon ceramic culture. According to the expression profile, only the administration of two factors (THPO and FLT3L) yielded in advanced cultivation conditions in the culture system establishment (Sieber et al., 2018). Furthermore, osteogenic differentiation induction and the generation of a 3D extracellular matrix network within the ceramic cavities might contribute to bone marrow niche mimicry. Also, differential expression of particular genes, that cannot be identified by the employed analysis approaches can be decisive in regulating the HSC pool in described bone marrow on a chip system. Notable, the transcriptome analysis was characterized by individual expression levels intrinsic to primary cells. Therefore, it is conceivable to apply the system for questions of personalized issues regarding HSC maintenance.

### Leaving the environment of the ceramic promotes differentiation

The significantly higher occurrence of native HSCs in the artificial *in vitro* bone marrow niche in the ceramic confirms the beneficial effect of the bone marrow niche on sustaining undifferentiated HSCs. The process of preserving the native state of the HSCs is a complex process in which various factors play a role. Besides others, it is an interplay of juxtacrine and paracrine signaling involving multiple cell types (Morrison and Scadden, 2014). In the circulation, the juxtacrine signaling between the stromal cells and the HSCs is not existent while the secreted signals are still present, explaining the reduced but not extinct native HSC population (Mayani, 2016). Furthermore, it remains unclear how long the native HSCs had resided in the circulation. It could be possible that the HSCs are continuously flushed out of the ceramic and subsequently start to differentiate when entering the circulation. This assumption is affirmed by including the data gathered for the expression of the differentiation markers CD36 and CD45RA. Cells differentiating towards the myeloid or erythroid lineage were found more abundantly in the circulation. Especially CD45RA expressing cells had a substantially higher occurrence in the circulation than in the ceramic. The reduction of native HSC during culture could be adjustment processes within the system. The amount of native HSCs is still too high compared to the *in vivo* situation where 0.01% are native HSCs (Brendel et al., 2014; Notta, F., Doulatov, S., Laurenti, E., Poeppl, A., Jurisica, I., and Dick, 2011). Additionally, the number of differentiated cells must increase to build up a functioning hematopoietic system.

### Rudimental granulopoiesis is observable within the MOC

Rudimental granulopoiesis towards the neutrophil lineage was observable in our model without the addition of further cytokines. Compared to their occurrence *in vivo* (0.9%) GMPs were more abundant in our model (~2-4%) while the incidence of myelocytes (~1-3%) was substantially smaller than *in vivo* (12.7%) (Greer, 1993). A significant number of neutrophils was not detectable. One reason might be a lack of G-CSF which is critical for neutrophil maturation. The introduction of endothelial cells, which are known to secrete G-CSF, could solve this problem. However, it might also lead to the mobilization of the HSCs (Garcia et al., 2015; Zhao et al., 2012).

### The model is robust

The removal of cells during medium exchange did not have a significant impact on the bone marrow niche. It is unknown if the cells, when removed, are flushed out of the ceramic or actively migrate. The former hypothesis appears more realistic since no vasculature is present within the model. Compared to the circulation, the ceramic CD34^+^CD38^−^ population only exhibited a small gradation between the 100%, 50%, and 0% values. This finding demonstrates the robustness of the bone marrow niche. The higher occurrence in the 0% reintroduction group in the circulation could mean that differentiated cells might have a negative influence on the sustainment of native HSCs.

Cell removal and subsequent analysis during a running experiment without destroying the bone marrow niche are possible. The intrinsic self-stabilization capacity provides the opportunity to continually monitor lineage changes or genotoxic effects in bone marrow safety studies without ending the experiment. In a larger scale, it could also be used to expand HSPCs prior implantation constantly.

### HSCs cultured in the bone marrow model on the MOC are capable of engrafting in irradiated immunocompromised mice

Transplantation of cells into irradiated immunocompromised mice remains the gold standard assay for the assessment of the repopulation capability of hematopoietic stem cells. The successful repopulation of a vacant bone marrow niche proves that the cells are native HSCs (Jingjing and ChengCheng, 2015; Nakamura-Ishizu et al., 2014).

Even though a significant difference in the T cell population measured after 12 weeks in the mouse, no other significant differences were observed in comparison to the control. Thus, the engraftment of the HSCs that were cultured for four weeks in the bone marrow model on the MOC was successful. Additionally, the sustained hematopoietic activity after 20 weeks proves that the transplanted cells were native HSCs capable of self-renewal and not MPPs which would have vanished after this time (Doulatov et al., 2012; Pineault and Abu-khader, 2015).

In a second experiment, HSCs were cultured for eight weeks in the bone marrow model on the MOC. The measured percentages were significantly lower compared to the control. However, all cell types were present even if monocytes and NK cells nearly vanished after 20 weeks. The low percentages might be ascribed to the small cell numbers of native HSCs (CD45RA^−^CD34^+^CD38^−^CD90^+^) which have been depicted in figure 8. Similar results have been described in the literature in cases where only a few native HSCs were used for engraftment experiments (Notta, F., Doulatov, S., Laurenti, E., Poeppl, A., Jurisica, I., and Dick, 2011).

As in the previous experiment, the engraftment was successful. A prolonged experiment time might have led to an increase in T cells. Nevertheless, all other cell types were present after 20 weeks of culture, again proving the existence of native HSC after an eight-week long culture of HSPCs in the bone marrow model on the MOC (Doulatov et al., 2012; Pineault and Abu-khader, 2015).

### Conclusion

The aim of this project was the generation of a model capable of long-term sustainment of primitive HSCs and its implementation on the MOC. In the first step, a suitable environment for long-term HSC culture was generated. In the second step, HSPCs isolated from umbilical cord blood were seeded in this bone marrow mimicking environment. It could be shown that HSCs remained their native phenotype for at least eight weeks in dynamic culture conditions. Furthermore, a significant deviation in the differentiation pattern was observed between HSPCs residing in the circulation and in the ceramic in the MOC. Significantly more native HSCs were present in the ceramic while significantly more differentiated HSPCs were identified in the circulation. Also, granulopoiesis could be observed without the addition of further cytokines.

In conclusion, the here presented bone marrow model demonstrates for the first time the successful long-term culture of functional multipotent HSCs in a dynamic environment. The model surpasses all previously presented 3D bone marrow models in culturing time of HSCs as well as in mimicking the *in vivo* environment (Kim et al., 2015). Predestining it as a model for sophisticated *in vitro* drug testing, thus, serving as an alternative to animal testing. It harbors the potential to be developed into a model mimicking the whole human bone marrow and in the farther future the entire hematopoietic system. The bone marrow model could be interconnected with other organs on an extended MOC laying the ground stone for the human on a chip.

## Material and Methods

### Isolation and expansion of MSCs

Human MSCs were isolated from the bone marrow of femoral heads as described in (Sieber et al., 2018). In brief, the cells were washed out, separated by density gradient centrifugation, and the PBMC layer seeded into a T25 culture flask. The cells were cultured in Dulbecco’s modified Eagle’s medium (DMEM) (Corning Inc., USA) + 10% Fetal Calf Serum (FCS) (Biochrom, Germany) + 1% Penicillin-Streptomycin (P/S) (Biowest, France) and used until passage 7.

### Isolation of HSPCs

Human HSPCs were isolated from umbilical cord blood as described in (Sieber et al., 2018) In brief, the blood was separated by density gradient centrifugation and HSPCs were segregated from the other cells using the Miltenyi MACS CD34^+^ isolation kit (Miltenyi Biotec, Germany). Cell number, phenotype, and purity were evaluated by flow cytometry according to CD34, CD38, CD45RA, CD49f, CD90, and CD133 (all Miltenyi Biotec, Germany) expression. The cells were resuspended in StemSpan ACF (Stemcell Technologies, USA) + 25 ng/ml FLT3-L + 10 ng/ml TPO (both PeproTech, USA) + 1% P/S.

### 3D cell culture

Hydroxyapatite-coated zirconium oxide based Sponceram 3D ceramic scaffolds (Zellwerk GmbH, Germany) were used as a scaffold for the 3D culture. The cylindrically shaped ceramics are 5.8 mm in height and diameter. The pore size in the ceramic is the same as observed for the human bone marrow. MSCs were seeded onto the scaffold in DMEM + 10% FCS + 1% P/S and cultured for seven days. Subsequently, medium was changed to Stemspan-ACF + 25 ng/ml FLT3-L + 10 ng/ml TPO + 1% P/S and 5,000 CD34^+^CD38^−^ HSPCs were added and allowed to adhere overnight in the incubator. The next morning, the ceramics were transferred to the MOC for dynamic culture. For medium exchange, the medium was collected and centrifuged at 300 g for 5 min. The supernatant was discarded, and the pellet resuspended in fresh medium and transferred back onto the ceramic. The medium was exchanged every 2 to 3 days.

In one experiment, ceramic and circulating HSPCs were analyzed separately, the ceramic was transferred from the culture compartment to a 24-well plate well for extraction of HSPCs. The remaining HSPCs in the MOC were defined as circulating cells.

In another experiment, after a two-week stabilization period, all, 50% or 0% of the HSPCs extracted during medium exchange were returned to the MOC. For the 50% return, half the medium was discarded before centrifugation. After centrifugation, the pellets were each resuspended in fresh medium and pipetted back into the medium or ceramic reservoir, respectively. For the 0% return, the whole medium cell suspension was discarded. The medium was exchanged for the last time as described above two days prior cells were being analyzed.

### Microfluidic system

The multi-organ-chip is a microfluidic platform which was developed in our institute. It consists of two separate independent circular channel systems. Each circuit is hosting two culture compartments interconnected by a channel system. The flow rate of the medium is controlled by a peristaltic on-chip micropump which is integrated into each circuit. One pump consists of two valves which surround one pumping membrane. All of them are operated by air pressure. The air pressure is generated by a control unit to which the MOC is connected via tubes. The control unit controls the flow rate and volume of the medium in the chip. The pump provides a pulsatile medium flow through 100 μm high and 500 μm wide channels. The pumping volume ranges 5-70 μl min^−1^ and the frequency 0.2-2.5 Hz. (Ataç et al., 2013; Marx et al., 2012; Maschmeyer et al., 2015; Sonntag et al., 2010). A pump frequency of 2 Hz for the continuous dynamic operation at a flow rate of 5 μl min^−1^ was used. The MOC was manufactured as described by Wagner et al. (Wagner et al., 2013). In brief, Wacker primer (Wacker Chemie, Germany) was applied to the adapter plate and incubated for 20 min at 80°C. Next, a casting chamber was prepared to consist of the prepared adapter plate, a master mold, and a casting frame. PDMS was injected into this casting station, and the whole setup was incubated for 60 min at 80°C. The resulting 2 mm thick PDMS layer containing the imprint of the pumps and channels was permanently bonded by low-pressure plasma oxidation to a glass slide with a footprint of 75 x 25 mm (Menzel, Germany), thereby forming the enclosed microfluidic channel system (Wagner et al., 2013).

The seeded ceramic was placed in one of the two culture compartments of each circuit. Only the outer compartments of the system were used for the ceramic while the inner served as medium reservoirs.

### Scanning electron microscopy (SEM)

Electron microscopy was performed in the department for electron microscopy (ZELMI) of the TU Berlin employing a Hitachi S-4000 (Hitachi, Japan). The bone marrow model was cultured for four weeks on the MOC. The medium was removed, the ceramics were carefully washed with PBS and afterward fixed in 4% PFA for three hours. The day before microscopy, samples were dried using an ascending sequence. The samples were mounted on a support using silver before being transferred to a vacuum chamber and sputter-coated with gold.

### Flow cytometry

Cells were stained with anti-human antibody for CD34-PE, CD38-APC, CD45RA-VioBlue, CD49f-FITC, CD90-FITC, CD133-FITC, CD36-FITC, CD15-PE-Vio770 and CD16-PerCP-Vio700 (all Miltenyi Biotec, Germany) in the dark for 10 min on ice. The analysis was carried out using a Miltenyi MACSQuant Analyzer flow cytometer (Miltenyi Biotec, Germany). Data were processed using FlowJo 10 (FlowJo LLC, USA).

### CFU-GEMM assay

The myeloid differentiation potential of the HSPCs extracted from the ceramic was assessed by colony forming unit - granulocyte, erythrocyte, macrophage, megakaryocyte (CFU-GEMM) assay using Miltenyi’s StemMACS HSC-CFU media (Miltenyi Biotec, Germany). The assay was performed following the instructions provided by the manufacturer.

### Transcriptome analysis via next generation sequencing

Transcriptome analysis was performed by using next-generation sequencing (NGS) from Illumina. First, a cDNA library was generated employing the TruSeq^®^ Stranded mRNA LT Sample Prep Kit (Illumina), according to the TruSeq^®^ Stranded mRNA Sample Preparation Protocol LS (Illumina). The initial quantity of total RNA was 800 ng for BMSCs and 50-110 ng for HSPCs. The cDNA was enriched via PCR. Diverse purification steps to separate the nucleic acid from reaction mix between the steps were performed with Mag-Bind^®^ RXNPure Plus magnetic beads (Omega Bio-Tec Inc.). As a quality control and to determine the concentration and size of the fragments, the cDNA was analyzed with a UV-Vis spectrophotometer (NanoDrop2000, Thermo Scientific) and gel electrophoresis (2% agarose).

The sequencing was carried out by the Illumina MiSeq^®^ System. Raw data, generated by the Illumina MiSeq^®^ platform, were processed on the Galaxy Project Platform (Afgan et al., 2018), using the tools FASTQ Groomer, to convert the output data into Sanger sequencing data. Fragments were then mapped against the human genome (hg38) to detect splice junctions between exons by HISAT2. Mapped reads were processed to the FeatureCount tool on the Galaxy Project Platform. Differential expression analysis was performed by the DESEQ2 algorithm (nominal p-value < 0,05; FD>2) provided by DE analysis (https://yanli.shinyapps.io/DEApp/ - App bioinformatics core, Center for Research Informatics (CRI), Biological Science Division (BSD), University of Chicago). Differentially regulated genes were analyzed for overrepresented gene sets by GSEA (Gene Set Enrichment Analysis – http://software.broadinstitute.org/gsea/index.jsp; broad institute, Cambridge;) (Subramanian, Tamayo, et al. (Subramanian et al., 2014). Heatmap generation was conducted by ClustVis analysis platform (Metsalu and Vilo, 2015) and the Morpheus tool from broad institute Cambridge (https://software.broadinstitute.org/morpheus).

### In vivo application of HSC into immunodeficient mice

For in vivo application of HSC NOD.cg-Prkdc scid Il2rg tm1 Saug (NOG) mouse strain from Taconic was used. Three-week-old NOG mice were irradiated with 1.5Gy at least 4h before HSC application.

HSC samples for direct application without in vitro culture were thawed in high percentage serum solution (50% FBS, 50%PBS). Cell number was determined, and 5x104 HSC were resuspended in 200μl per mice for intravenous application. HSC samples from in vitro expansion were also counted, and the same cell number was adjusted. The whole 200 μl cell suspension was subsequently injected into the tail vein of the mice.

Blood samples were taken after 4, 8, 12, 16, 20 and 24 weeks. Blood cells were analyzed using flow cytometry. After four and eight weeks, the cells were stained for CD45 and HLA (Miltenyi). At all other points in time, the cells were stained with the 7-Color-Immunophenotyping Kit from Miltenyi. Samples were blocked with human and murine FcR blocking reagent and antibodies were used concerning manufactures instructions. Blood lysis was employed at the end of the staining protocol to eliminate erythrocytes from samples.

### Statistical analysis

Unpaired t-test was applied to the data sets, using GraphPad Prism software version 6.04 (GraphPad Software Inc., USA). P values smaller than or equal to 0.05 were considered significant.

## Acknowledgements

The authors sincerely thank the Vivantes Hospital in Friedrichshain, Berlin for providing umbilical cord blood samples and the Immanuel Hospital in Berlin for providing bone marrow samples. We also thank Alexandra Lorenz for designing the bone marrow culture compartments. Contributions by Stefan Sieber were made possible by DFG funding through the Berlin-Brandenburg School for Regenerative Therapies GSC 203. The work has been funded by the German Federal Ministry for Education and Research, ERA-Net “EuroTransBio”, Grant No.031A597B.

